# Enhancement of RecET-mediated *in vivo* linear DNA assembly by a *xonA* mutation

**DOI:** 10.1101/2022.01.13.476211

**Authors:** James A. Sawitzke, Nina Costantino, Ellen Hutchinson, Lynn C. Thomason, Donald L. Court

## Abstract

Assembly of intact, replicating plasmids from linear DNA fragments introduced into bacterial cells, i.e. *in vivo* cloning, is a facile genetic engineering technology that avoids many of the problems associated with standard *in vitro* cloning. Here we report characterization of various parameters of *in vivo* linear DNA assembly mediated by either the RecET recombination system or the bacteriophage λ Red recombination system. As previously observed, RecET is superior to Red for this reaction when the terminal homology is 50 bases. Deletion of the *E. coli xonA* gene, encoding Exonuclease I, a 3’→5’ single-strand DNA exonuclease, substantially improves the efficiency of *in vivo* linear DNA assembly for both systems. Deletion of ExoI function allowed robust RecET assembly of six DNA segments to create a functional plasmid. The linear DNAs are joined accurately with very few errors. This discovery provides a significant improvement to previously reported *in vivo* linear DNA assembly technologies.

## Introduction

Recombineering is *in vivo* genetic engineering using bacteriophage-encoded recombination proteins able to act on short (~50 base) homologies. These proteins promote recombination between the bacterial chromosome or episomes and linear DNA, either double-stranded (dsDNA)^1, 2^ or single-stranded (ssDNA)^3^. Such phage systems typically include two proteins that are expressed and act in a coordinated manner^4^. One protein, a 5’→3’ dsDNA-dependent exonuclease^5^, processes linear dsDNA to leave single-strand 3’ overhangs that are bound by the second protein, a single-strand annealing protein (SSAP)^6^. This SSAP-bound ssDNA is annealed to its complementary single-strand sequence in the bacteria. Both proteins are required for recombination of dsDNA^1, 2, 6, 7^, while only the SSAP is required for ssDNA recombination^3^. In phage λ these proteins are the exonuclease, Exo^5^, and the SSAP, Beta^6–8^. A second commonly used recombineering system, RecET, is derived from the *E. coli* cryptic prophage Rac^9^. The RecE protein is a 5’→3’ dsDNA-dependent exonuclease^10^, however, RecE (866aa) is nearly four times the size of λ Exo (226aa), with the exonuclease domain contained in the last ~260aa^10, 11^. The function of the large, dispensable N-terminus of RecE is unknown. RecT (269aa), like λ Beta (261aa), is a SSAP^12^. Phage λ also encodes an inhibitor of the *E. coli* RecBCD nuclease, the Gam protein^13^, which preserves linear dsDNA in the bacterial cell. The Rac prophage does not contain a *gam* gene, although it has been suggested that an equivalent inhibitory function is encoded within the large *recE* gene^14^. λ *gam* has been included in some RecET expression constructs and increases RecET-mediated recombination frequencies in some assays^14, 15^. While RecE and RecT play analogous roles to λ Exo and Beta^1, 12^, and RecET can replace Exo and Beta for λ growth and recombination^12, 16^, there is little sequence identity between the two systems other than a few key amino acids^17, 18^. Thus, while both systems can be used in *E. coli* for recombineering, they may differ in their mechanistic details.

A useful genetic engineering technique is the homology-dependent *in vivo* assembly of multiple linear dsDNAs into a single DNA molecule, usually a plasmid^15^. The linear DNAs containing the required homologies can be introduced into recombination-proficient cells by transformation^19^. Both the Red and RecET recombination systems will perform this homology-dependent reaction, but the RecET system gives higher recombination effiencies than does Red^15^.

Linear DNA assembly can also be achieved in DH5α^20–22^, a commercially available strain commonly used for plasmid propagation and cloning; or in JC8679, a strain that expresses RecET^23^. Nozaki and Niki^21^ have studied and optimized linear DNA assembly in DH5α and other *E. coli* strains. Exonuclease III function is required; ExoIII is a processive 3’→5’ exonuclease encoded by the *xthA* gene^24^. Processing of linear DNA by this nuclease leaves 5’ overhangs, of opposite polarity to the 3’ overhangs formed by the phage exonucleases. They also found that mutation of the host specificity restriction system (*hsdR*) was important for obtaining high recombination efficiencies^21^.

*In vivo* cloning has been adapted for bioprospecting^15^ and complex plasmid constructions. This method allows simultaneous incorporation of several genetic elements, such as promoters and gene tags. The technique avoids the difficulty of sequentially cloning these individual elements and can provide a simple and inexpensive *in vivo* alternative to Gibson Assembly^25^. In this paper we report characterization of various parameters of the *in vivo* linear DNA assembly reaction mediated by RecET and λ Red. With both systems, we find that deletion of the *E. coli xonA* gene, encoding Exonuclease I, a 3’→5’ ssDNA exonuclease^26^, improves the efficiency of this DNA assembly reaction. For the RecET system it allows robust assembly of at least six DNA fragments in a single reaction. The linear DNAs are joined accurately with very few errors. This discovery is a substantial improvement of *in vivo* linear DNA assembly as compared to other published systems^15, 20–23, 27–30^

## Results and Discussion

### *In vivo* assembly of two linear dsDNAs

We first confirmed published observations^15^ using the Red and RecET systems. Initially, strains expressing either the λ Red or RecET functions were compared for their ability to assemble plasmids *in vivo* from two linear DNAs introduced into cells by electroporation. Recombination functions were expressed from the λ *P*_L_ promoter in single copy on the bacterial chromosome, under control of the temperature sensitive CI857 repressor (Figure 1A) and standard recombineering techniques were used^19^. One strain contains the phage λ Red system (HME6), and another contains the RecET system (LT1795); both strains express λ *gam*. Strains lacking *gam*, λ Red (JS663) and RecET (SIMD63), were also tested.

**Figure 1.**
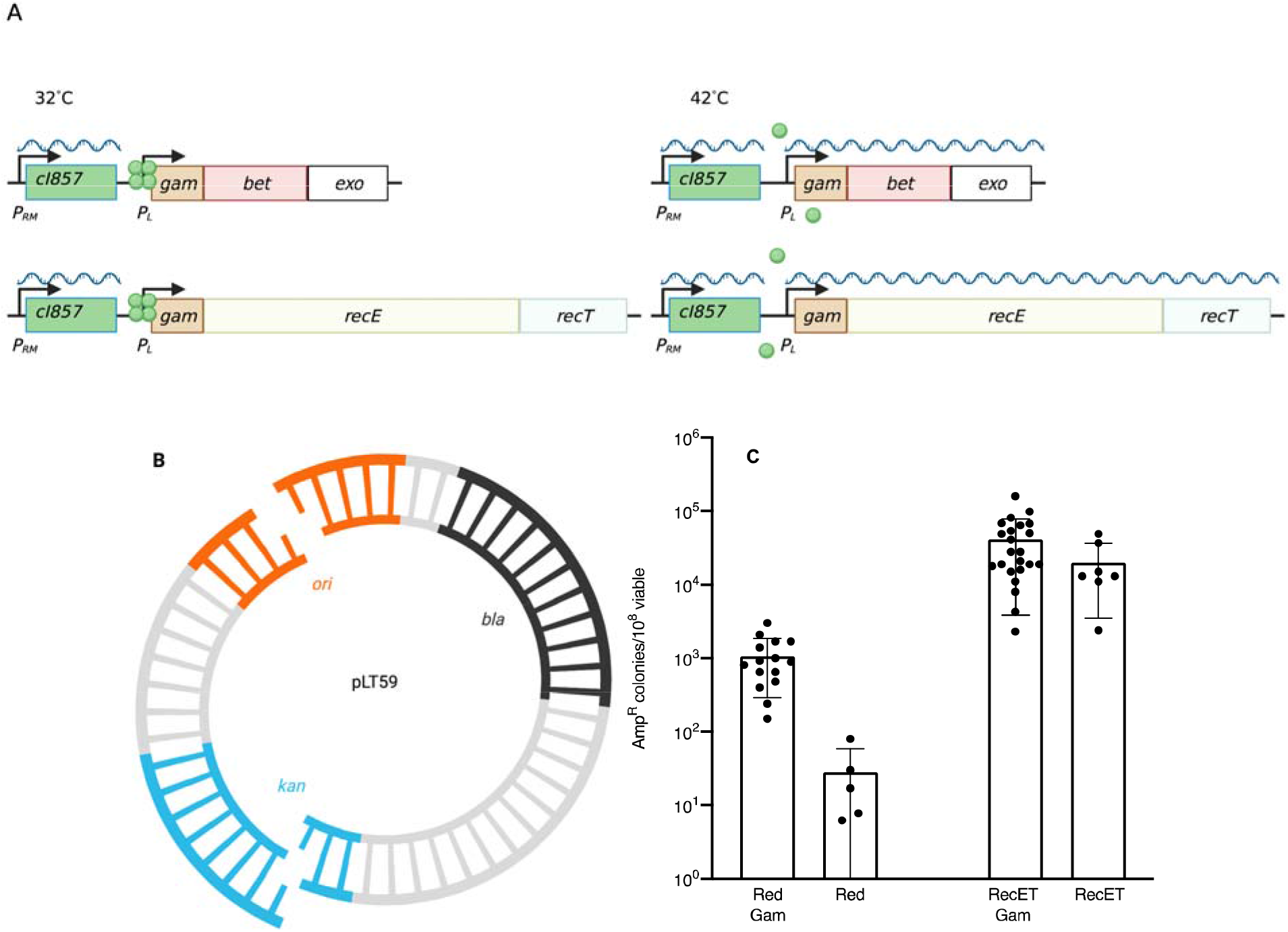
Assembly of two linear DNAs into intact plasmids by λ Red and RecET systems. **A.** Expression of Red or RecET from the *P*_L_ operon of a defective λ prophage. At 32°C, the *C*I857 repressor protein (green circle) is functional and the *red* or *recET* genes are not transcribed. At 42°C, the CI857 repressor unfolds and no longer binds to the operators allowing transcription of the recombination genes from the *P_L_* promoter. **B.** Two-fragment linear DNA assembly. 50bp of terminal homology is present in each fragment within the *ori* (origin of DNA replication) or the *kan* gene, indicated as the single-strand bases. *bla* encodes β-lactamase which when expressed results in ampicillin resistance (Amp^R^) **C.** Recombinant frequency is expressed as the number of Amp^R^ colonies/10^8^ total colonies for the indicated recombineering systems. Error bars indicate standand deviation (s.d.). Results from independent experiments are indicated by the black dots. Assembly was dependent on supplying both DNA fragments and on expression of the λ Red or RecET functions.

Two dsDNA fragments were used to generate an intact plasmid (pLT59). Each DNA fragment contained a partial origin of DNA replication (*ori*) and a partial kanamycin resistance (*kan*) gene (Figure 1B). Intact plasmids were selected by resistance to ampicillin (Amp^R^). Only precise joining of the two DNAs will generate a functional plasmid *ori* and a Kan^R^ gene. The results are shown in Figure 1.

As previously reported^15^ the RecET system is up to 1000-fold more efficient at *in vivo* linear DNA assembly than is the λ Red system (Figure 1C, Red vs. RecET). However, we found that increasing the length of terminal homology increased the frequency of λ Red-mediated recombination such that with >330bp of homology the difference between the two systems is only 5-fold (Supporting Information, Figure S1). This difference in homology length requirements suggests interesting mechanistic differences between the systems that are beyond the scope of this letter. When Amp^R^ colonies were patched to LB agar containing kanamycin the frequency of Kan^R^ was >95% for all strains tested, indicating accurate joining of the junction within the *kan* gene by either recombination system. Expression of Gam stimulated Red-dependent recombination and plasmid assembly ~10-fold, as expected from previous recombination experiments^31^. Interestingly, the presence or absence of Gam did not alter the frequency of linear DNA assembly by RecET, in agreement with a proposal^14^ that the larger RecE protein contains a Gam-like activity. However, from an abundance of caution, in all other RecET experiments described here we used the Gam-expressing strains to ensure protection of linear DNA from RecBCD.

### Mutation of Exonuclease I function improves recombinant yield for linear DNA assembly

Processing of linear dsDNA ends by either λ Exo or RecE leaves 3’ ssDNA overhangs. If not bound and protected by Beta or RecT, these single strands could be substrates for bacterial 3’→5’ exonucleases. Thus, degradation by a host 3’→5’ exonuclease could lower recombination efficiency by removing DNA substrate. Since ExoI is a major *E. coli* 3’→5’ ssDNA exonuclease^26^, we asked whether removing ExoI function by deletion of the *xonA* gene improves the efficiency of linear DNA assembly. We tested the recombination proficiency of strains expressing either λ Red (NC540) or RecET (NC553) deleted for the *xonA* gene (Δ*xonA*). As shown in Figure 2, removal of ExoI activity enhanced recovery of recombinant plasmids nearly ~65-fold for the Red system and >70-fold for RecET. Amp^R^ colonies from the strains deleted for *xonA* were patched to L+Kan; >97% of the isolates were Kan^R^, indicating the loss of ExoI function did not affect fidelity of fragment joining.

**Figure 2.**
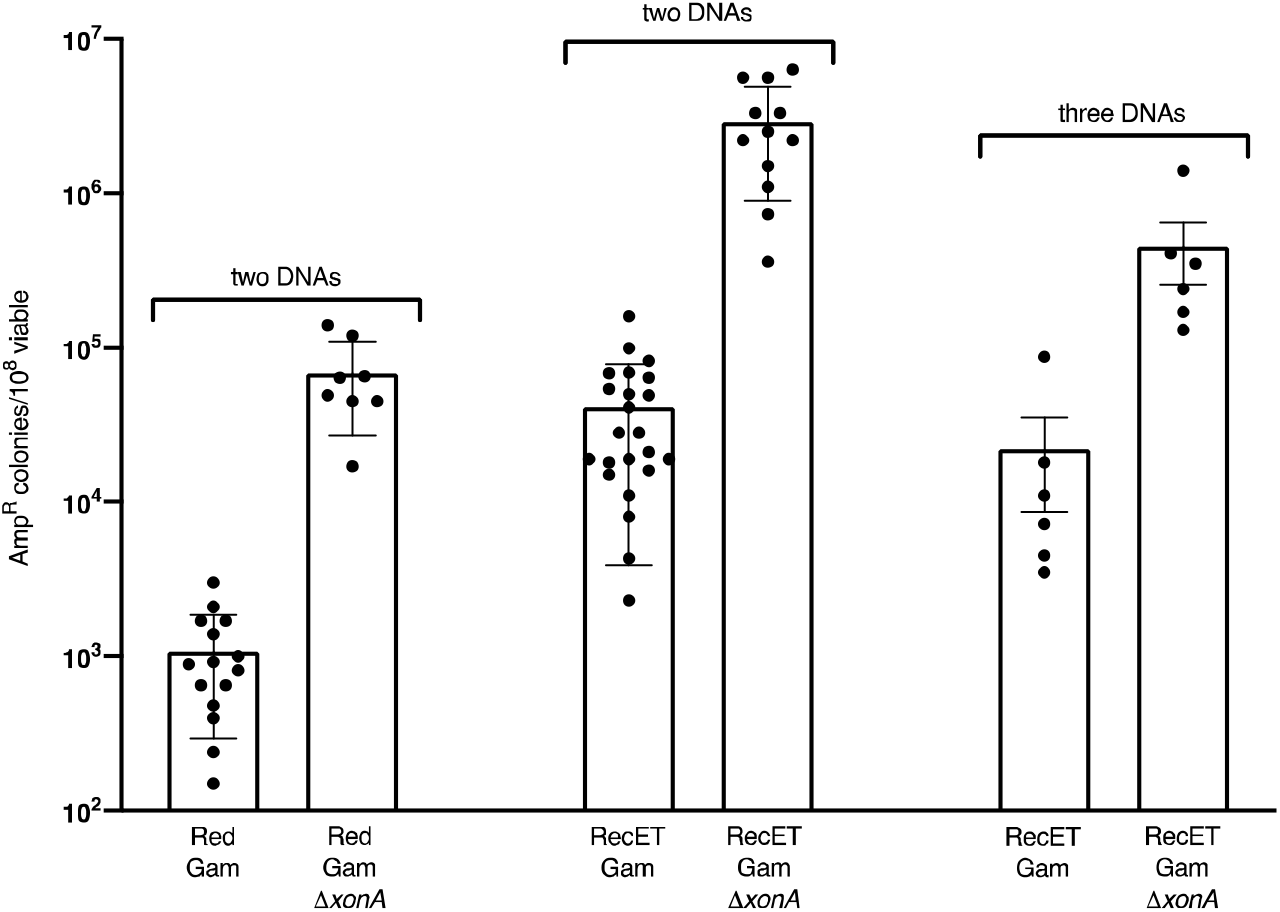
Removal of Exonuclease I activity improves recombinant yield. The recombinant yield is expressed as the number of Amp^R^ colonies/10^8^ total colonies for the Red and RecET systems, both with Gam expressed. Results from independent experiments are indicated by the black dots. The s.d. is shown for two fragment assembly and standard error of the mean (s.e.m.) for three fragment assembly

We also tested the ability of the RecET strain to assemble three fragments to make the same intact plasmid (pLT59) (Figure S2). As before, the DNA junctions bisected *ori* and the *kan* gene, but in this case the *bla* gene encoding ampicillin resistance was also split into two separate DNA fragments so that accurate assembly was necessary for plasmid replication and resistance to ampicillin. The three fragments were efficiently assembled by the RecET system, and elimination of the ExoI function increased the assembly frequency by ~20-fold.

### Two fragment assembly: dependence on DNA concentration

Since the RecET system is more efficient than the λ Red system for linear DNA assembly, we examined other aspects of RecET-mediated recombination. To ask whether recombination efficiency changes with DNA concentration, a range of linear DNA concentrations was tested in both the *xonA^+^* and Δ*xonA* RecET expressing strains (Figure 3). The Δ*xonA* strain showed robust levels of fragment joining even at the lowest DNA concentration tested, 10ng per fragment.

**Figure 3.**
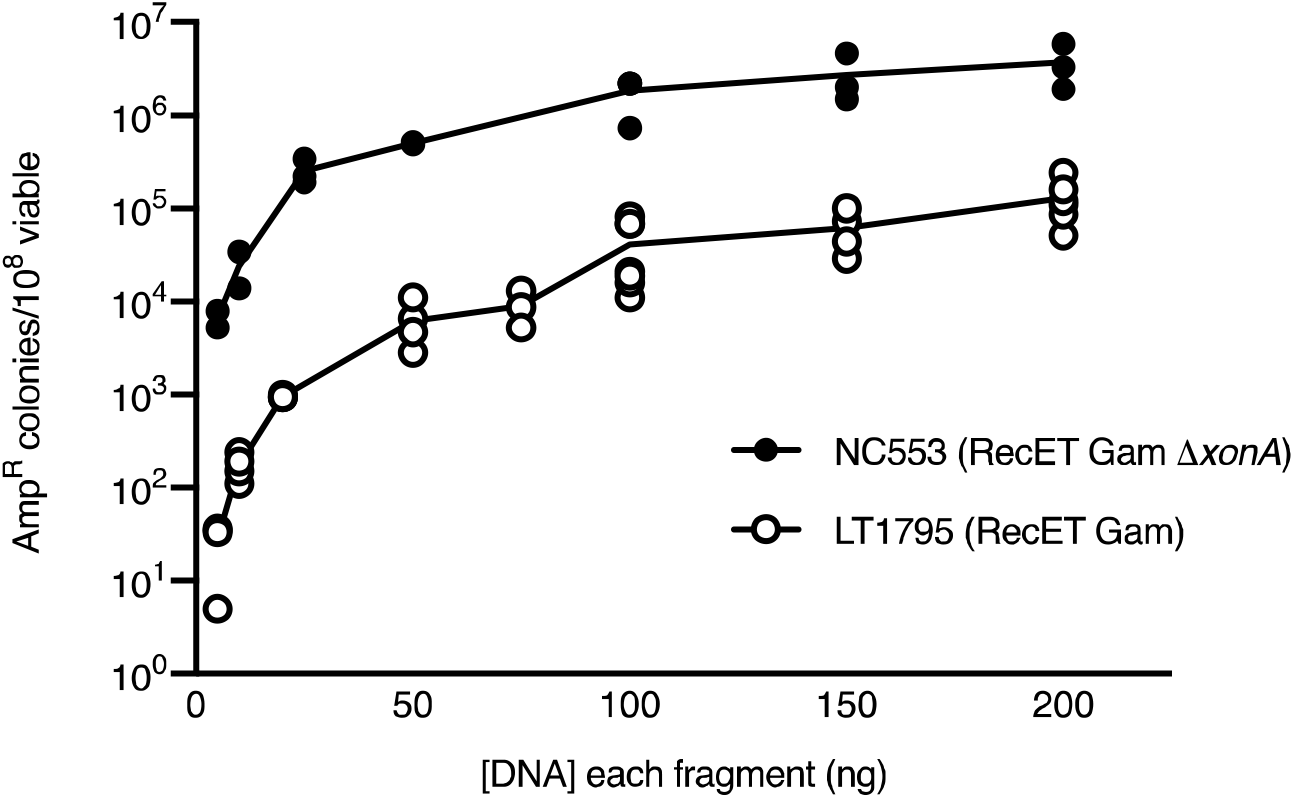
Dependence of two-fragment linear DNA assembly on DNA concentration. The data show number of Amp^R^ recombinants/10^8^ viable cells over a range of DNA concentrations. The concentration of each linear dsDNA is indicated. Results from independent experiments are indicated by the circles. Closed circles (•) indicate RecET Δ*xonA* (NC553); open circles (⍰) indicate RecET *xonA^+^*(LT1795).

### Two fragment assembly: recombination dependence on homology length

We asked whether shorter homology lengths would be efficient for fragment joining. We used a similar design to assemble two linear DNAs as shown in Figure 1B. Using PCR, we amplified pairs of linear dsDNAs with terminal homologies of various lengths, and tested these fragments in the RecET *xonA^+^* vs. the Δ*xonA* strain. As shown in Figure 4, 30bp of terminal homology provides nearly maximal recombination capability, similar to results seen previously in DH5α^21, 22, 27^. Our RecET results differ from those of Fu *et al.*^15^ (Supporting Information, Fig S1), who found few recombinants using 30bp homologies in a RecET-expressing strain retaining ExoI activity although they observed an increase in recombinant frequency as homology length increased from 50bp to 120bp. Because longer homologies are not required for efficient fragment assembly in the RecET Δ*xonA* strain, PCR products can be generated using DNA oligonucleotides (oligos) as short as 50nt in total length (30nt homology plus 20nt for priming).

**Figure 4.**
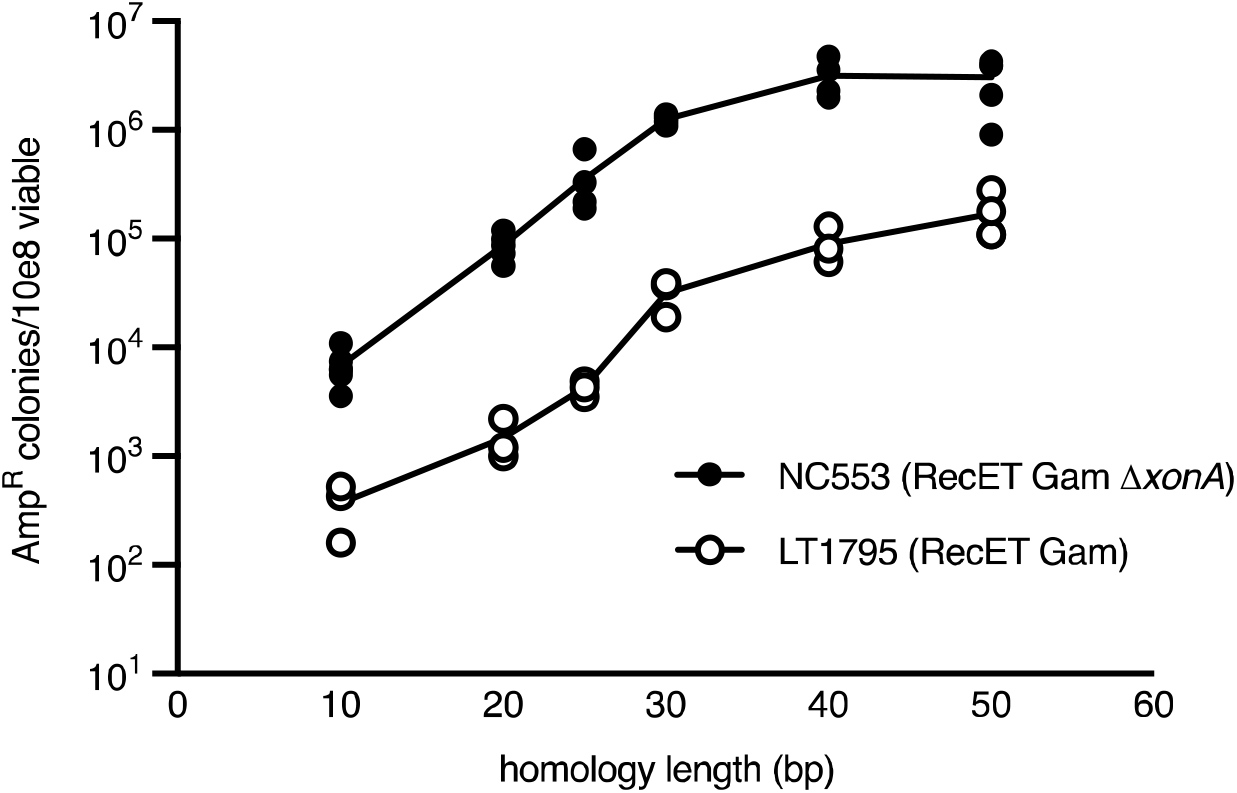
Dependence of two-fragment linear DNA assembly on terminal homology length. The data show the number of Amp^R^ recombinant plasmids/10^8^ viable cells obtained over a range of homology lengths. Results from independent experiments are indicated by the circles. Closed circles (•) indicate RecET Δ*xonA* (NC553); open circles (○) indicate RecET *xonA^+^*(LT1795).

### *In vivo* linear dsDNA assembly of plasmids from six linear dsDNA fragments

To test assembly of six DNA fragments, we designed a plasmid, pBR-*lacZ*, with the *E. coli lacZ* open reading frame replacing the tetracycline resistance gene, *tet*, of pBR322 (Figure 5A). The three fragments comprising the *lacZ* gene were either made by PCR or synthesized as gBlocks. The plasmid backbone was contained in three PCR fragments. After the recombination reaction, cells were plated on LB+Amp+X-gal to screen for blue colonies indicating β-galactosidase activity and thus accurate assembly of the *lacZ* gene. When six PCR fragments were used to assemble pBR-*lacZ* in the Δ*xonA* strain expressing RecET, the recombinant frequency was nearly 4.0×10^4^/10^8^ viable cells (Figure 5B), only ~15-fold lower than the frequency of the three-way reaction in the Δ*xonA* background (see Figure 2). Reducing the DNA concentration of each fragment from 100ng to 33ng may be responsible for much of this effect (Figure 3). When all fragments were generated with PCR, ~90% of the Amp^R^ colonies obtained from the Δ*xonA* strain were blue on LB+Amp+X-gal indicator agar, demonstrating β-galactosidase activity and thus correct assembly of the *lacZ*^+^gene. When the three fragments encoding *lacZ* were synthesized as gBlocks (IDT), 84% of the colonies were blue on LB+Amp+X-gal indicator agar. We isolated a total of 113 plasmids from independent white colonies for either the PCR or PCR/gBlock reactions performed in the Δ*xonA* background. These plasmids were analyzed by restriction analysis (Materials and Methods); in all but one case the six fragments had assembled correctly. Forty of the 113 plasmids isolated from white colonies were further examined by sequencing the *lacZ* gene and flanking region; this analysis revealed that most mistakes arise during PCR or gBlock synthesis rather than occurring at the recombinant junction (see Figure S3). In contrast, for LT1795, the *xonA*^+^ strain, increasing the number of linear DNA fragments from three to six caused a ~340-fold reduction in frequency, with only ~25% of the colonies blue on X-gal indicator. Restriction analysis of four white colonies from this strain revealed incorrect plasmid assembly.

**Figure 5.**
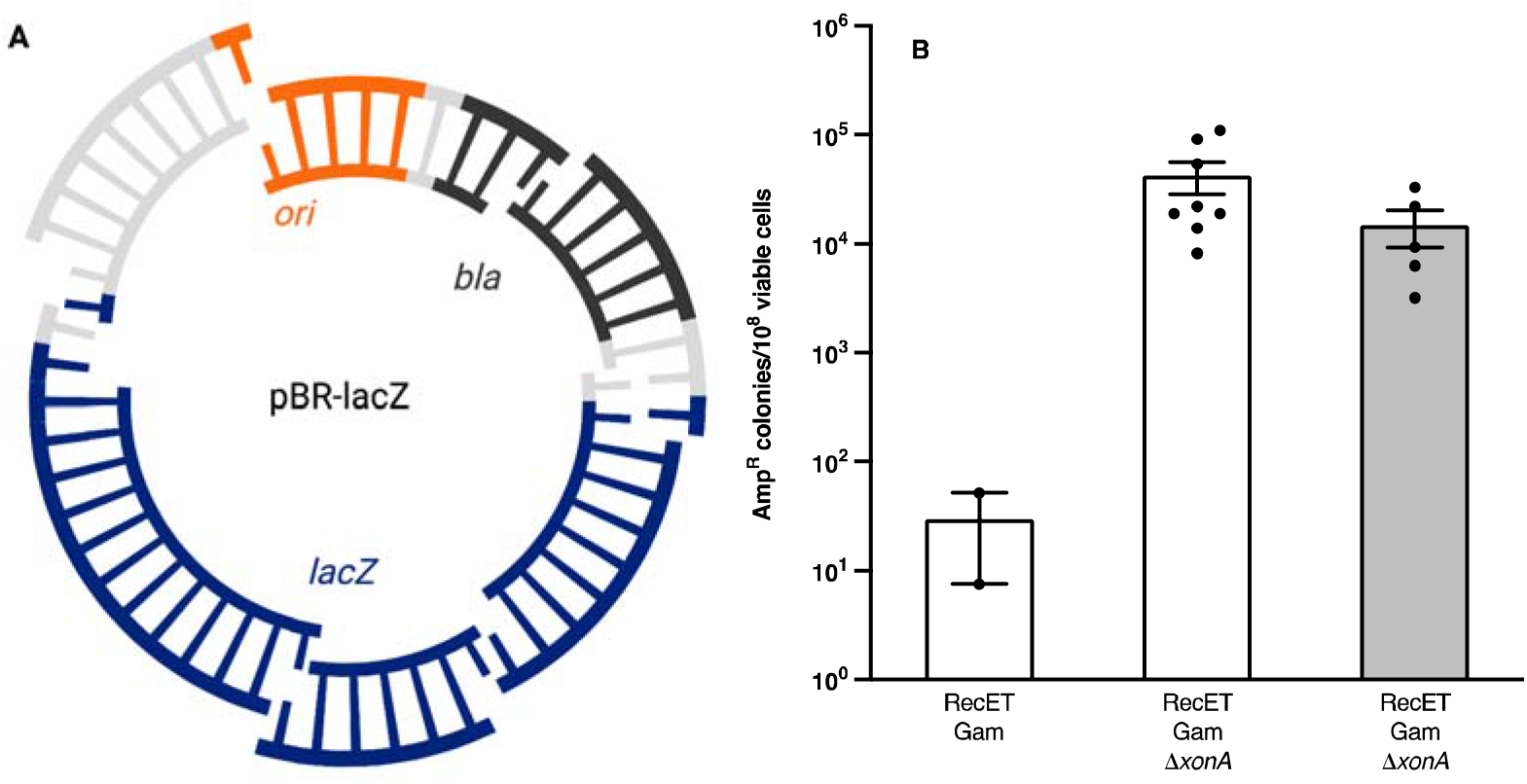
Assembly of six linear DNA fragments by RecET. **A.** Diagram of six-way plasmid assembly for pBR-*lacZ*. Terminal homologies present in each fragment within the *ori*, *bla*, *cat* and *lacZ* genes are indicated by the single-strand bases. One single-strand base indicates a homology of 50-60bp whereas two single-strand bases indicates a homology of ~100bp. Similar results were seen when all homologies were 50bp (Figure S4). **B.** Recombinant frequency obtained for six-way plasmid assembly under *xonA^+^* (LT1795) and Δ*xonA* (NC553) conditions is expressed as the number of Amp^R^ recombinant plasmids/10^8^ viable cells. All experiments used linear dsDNA generated with PCR except for that shown with the gray bar, where the three linear DNAs comprising the plasmid backbone were made with PCR and the three comprising the *lacZ* gene were synthesized as gBlocks. Results from independent experiments are indicated by the black dots. Error bars indicate s.e.m.

Our results (Figure 5B) show that elimination of the host ExoI function by deletion of the *xonA* gene substantially increases the efficiency of linear DNA assembly, allowing at least six fragments to be assembled by the RecET system into a functional plasmid (see Figure S4 of Supporting Information for additional data). Using the rigorous test of assembling six linear DNA fragments, we asked whether removal of other host ssDNA exonucleases impacted plasmid assembly. Removal of either the RecJ or ExoX functions did not increase recombination frequencies in the way that removing ExoI did (Supporting Information).

Several publications have demonstrated the power of *in vivo* cloning using phage recombination systems^15, 23, 27–30^ or an endogenous recombination activity present in some laboratory bacterial strains (DH5α)^20–22, 28^. This method of plasmid assembly allows complex plasmids to be designed and constructed with precision and accuracy. Regulatory elements, gene tags, fusions, and antibiotic resistance genes can all be introduced onto a plasmid backbone in a single round of *in vivo* cloning. However, as the number of fragments to be assembled increases, cloning efficiency rapidly decreases.^20, 21, 27, 28^ In an attempt to compare recombination efficiencies for various numbers of fragments, we have compiled a table (Supporting Information Table S1) of previously reported numbers of recombinants obtained by several laboratories and have compared that data with our results using the RecET system in the Δ*xonA* background. Although this comparison does have limitations since the methods used in the various laboratories were not uniform, the results are informative and are presented in graphical form in Figure 6.

**Figure 6.**
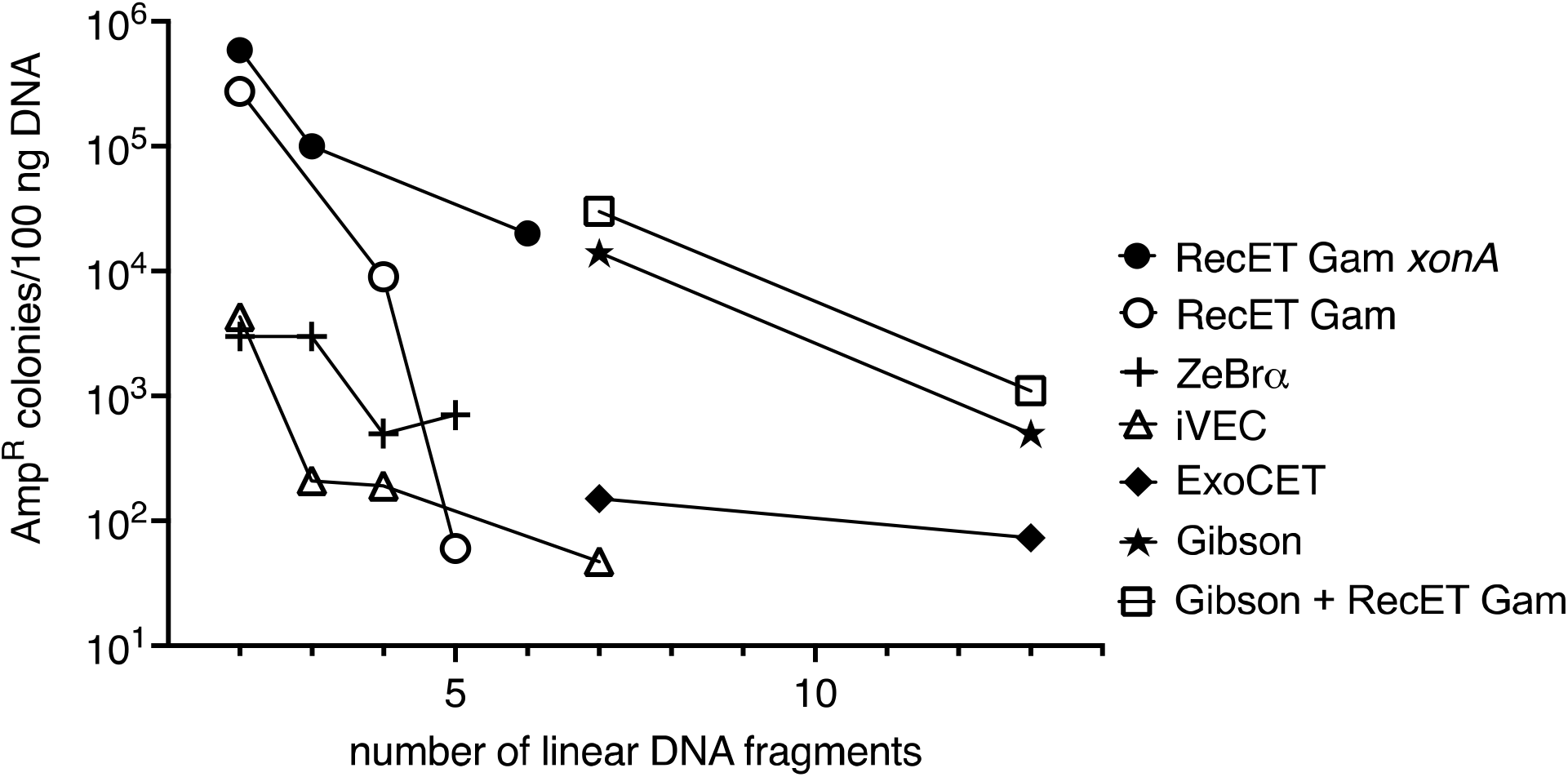
Comparison of numbers of recombinant plasmids generated by various *in vivo* cloning methods. The numbers of drug resistant plasmids obtained with the various methods of *in vivo* plasmid assembly are compared. Reported values were normalized to 100ng of input DNA per fragment. See Table S1 in Supporting Information for details. In all cases, each point represents the average value of multiple experiments. Shown are the data reported here for RecET Gam Δ*xonA* (filled circles), and published results for RecET Gam^15, 29^ (open circles), iVEC^21^ (open triangles), ZeBrα^28^ (plus signs), Gibson+RecET Gam^30^ (open squares), Gibson^30^ (stars), and ExoCET^30^ (filled diamonds).

It can be seen from the graph that the RecET Δ*xonA* strain (•) gives robust numbers of recombinant plasmids even when six fragments are assembled. These values are similar to those obtained by Gibson Assembly or Gibson Assembly + ExoCET for seven fragments^30^, both of which have lower accurancy than RecET Δ*xonA*^30^. The data of Fu *et al.*^15^ and Baker *et al.*^29^ (open circles), also show robust results for two and four fragments but fall off sharply when assembly of five different fragments is attempted. It should be noted that Fu *et al.*^15^ often express RecET from a plasmid rather than the *E. coli* chromosome, and when they assembled more fragments, they found four-fold lower values when the *recET* genes were expressed from the *E. coli* chromosome^29^. We note that for obvious reasons, it is preferable to avoid having a resident plasmid for recombinase expression in the same cells used for *in vivo* linear DNA assembly to generate another plasmid.

For the other two systems compared here, Nozaki and Niki^21^ successfully assembled seven fragments in their optimized iVEC strain, but obtained only a total of 40 colonies, suggesting that in this background, the seven-fragment assembly reaction is approaching the limits of recoverability. Richter^28^ reported assembly of linear DNA fragments *in vitro* using cell extracts containing the λ Red proteins (or not) and a specialized suicide vector; this mixture was purified and then transformed into NEB5α (i.e. DH5α). They found a significant decrease in assembly of four DNA fragments compared to three, but found similar frequencies with either four or five fragments (see Figure 6).

ExoCET cloning requires *in vitro* treatment of the DNA with T4 polymerase and was developed to increase the frequency of “direct DNA cloning”, i.e. retrieval of a specific region of DNA from a complex genome^32^. This method allows retrieval from a genomic mix at a frequency of up to 1.6×10^4^ (normalized to 100ng vector). When ExoCET was used to assemble multiple fragments directly (not a complex genome mix) and the results compared to those obtained with Gibson Assembly, Song *et al*.^30^ showed that ExoCET was superior to Gibson Assembly for obtaining correct clones contaning all fragments. Seven fragments were assembled at a frequency of 1.5×10^2^ (normalized to 100ng vector); and even 13 fragments could be assembled at a frequency of 7.3×10^1^. Our experiments raise the interesting question as to whether a *xonA* mutation would enhance the efficiency of the ExoCET reaction and perhaps direct cloning. We do not believe that Δ*xonA* would increase the frequencies seen in DH5α^21^ because the exposed ssDNAs for that system have a 5’→3’polarity.

Our data demonstrates that use of the RecET Δ*xonA* strain gives high levels of accurate recombination using nanogram quantities of DNA and is likely to be the method of choice for *in vivo* assembly of more than two or three fragments simultaneously. Additionally, the technique does not require special vectors, extract preparation, or *in vitro* reactions before transformation into competent cells. It also outperforms Gibson assembly, which is prone to omit fragments^30^, which we have not observed with RecET Δ*xonA*. We have used the RecET Δ*xonA* strain to make plasmids for a two-hybrid analysis by recombining a linearized vector with a PCR-generated DNA^33^. The high efficiency of our system allows plentiful recovery of recombinants for analysis. Recombinant clones can be identified either by a colony PCR screen or by plasmid DNA isolation and analysis (Supporting Information). Like others^15^, we have observed that for RecET the homology need not be exactly at the end of the linear DNA and that terminal non-homologies (from a few bases to >5 kb) will be removed during the recombination process, unlike ExoCET recombination^30^. Although we have not tested assembling more than six DNA fragments into a single plasmid, we predict that it should be possible in the RecET Δ*xonA* strain. We propose that the Δ*xonA* mutation allows the 3’ ssDNA ends produced by RecE to persist in the cell and thus allow enhanced RecT-dependent annealing of complementary sequences to form recombinant products.

## Materials and Methods

### Strains, plasmids and growth conditions

Bacterial strains are derivatives of *E. coli* K-12 (Table S2). LB media, both liquid and solid agar plates were used for growth. Ampicillin was used at 100 μg/ml. To test antibiotic resistance of recombinant plasmids, Amp^R^ colonies were patched to LB plates containing either kanamycin at 30 μg/ml or chloramphenicol at 10 μg/ml. To screen for blue colonies indicative of correct *lacZ* assembly and β-galactosidase activity, X-gal was added to LB solid agar plates at a final concentration of 50 μg/ml.

### Generation of linear fragments

Linear DNA fragments were generated by PCR using Platinum Taq HiFi (Invitrogen) and purified using a QIAquick PCR Purification kit (Qiagen). DNA oligonucleotides and gBlocks were obtained from IDT (https://www.idtdna.com). Sequences of DNA oligos are available upon request. Fragments contained 50bp of terminal homology unless otherwise noted. Fragments were generated from linear templates as described in Supporting Information, except for *lacZ* fragments, which were generated by colony PCR^19^. Purified linear DNAs were introduced individually into the relevant host by electroporation and ampicillin selection applied to ensure absence of intact plasmid contamination. In fragment assembly experiments, 100ng of each linear DNA was used unless noted. To prevent arcing during electroporation, in six-fragment assembly assays, equal amounts of individual fragments at 100ng/ul were mixed, keeping the overall DNA concentration at 100ng/ul but each individual fragment at 16.7ng/ul. Cells were electroporated with 2ul of this mixture, thus, the final concentration of each fragment was 33ng/ul.

### Expression of Recombination Functions and Electroporation

Cells were induced for the recombination functions (Figure 1A) and prepared for electroporation as previously described^19^. Following introduction of the DNA by electroporation, cells were outgrown in 1ml LB for 2 hrs at 32°C, diluted and plated on solid LB medium to score total viable cells, and on LB+Amp to score Amp^R^ plasmid recombinants. Petri plates were incubated at 32°C. In some experiments Amp^R^ colonies were patched to either LB+Kan or LB+Cm to screen for resistance. The frequency of fragment joining is calculated as (Amp^R^/Viable cells)(1×10^8^) since 1×10^8^ is approximately the number of cells that survive an electroporation under our conditions.

### Analysis of pBR-lacZ plasmids from transformants

In select experiments, white (Lac^−^) Amp^R^ colonies were purified on LB+Amp+X-gal, single colonies were used to grow 5ml overnight cultures in LB+Amp broth, and plasmid DNA was isolated using a Qiagen miniprep kit. Plasmids were digested with *Pst*I, which has a single restriction site in the *bla* gene and DNA was analyzed on agarose gels to verify that the recovered plasmids were of the expected size. Some plasmids were further analyzed by sequencing the *lacZ* gene.

## Supporting Information

Supplemental Results and Discussion

Effect of increasing terminal homology length on linear DNA assembly

Effect of *recJ* and *exoX* mutations on linear DNA assembly

Supplemental Methods for experiments described in this paper

Preparing linear DNA molecules for use in assembly

General procedures for linear DNA assembly

Table S1. Comparison of values for *in vivo* linear DNA assembly reported by different labs

Table S2. *Escherichia coli* K-12 strains and plasmids

Table S3. Supplemental *Escherichia coli* K-12 strains

Figure S1. Assembly of two linear DNAs with various homology lengths into intact plasmids by the λ Red and RecET systems

Figure S2. DNA fragments used for the three-way assembly of pLT59

Figure S3. Sequence analysis of white colonies from the pBR-*lacZ* assembly reactions

Figure S4. Additional data for *in vivo* linear assembly from six linear dsDNA fragments Supplemental References

## Acknowledgements

We thank Carolyn Court for careful reading of the manuscript and insightful comments. Comments from two anonomous reviewers improved the paper. We also thank M. Spencer, N. Shrader, T. Hartley, and K. Pike from the CRTP Genomics Laboratory at the Frederick National Lab for Sanger sequencing. All images in figures were created in their entirety by the authors as follows: Figures 1A, 1B, 5A, S2, and S4A were created with BioRender.com; Figures 1C, 2, 3, 4, 5B. 6, S1, S2, and S4B were made with Graphpad Prism; and Figure S3 was generated using Snapgene. This work was supported, in part, by the Intramural Research Program of the National Institutes of Health, National Cancer Institute, Center for Cancer Research. This project has also been partly funded with federal funds from the National Cancer Institute, National Institutes of Health, under contract no. HHSN261200800001E. Funding was provided by EMBL to J.A.S. The authors declare no conflicts of interest.

## Author Contributions

J.A.S. performed experiments, project design, wrote paper; N.C. performed experiments, project design, wrote paper; L.C.T. project design, wrote paper; E.H. performed experiments; D.L.C. project design, provided funding.

---

For Table of Contents Only

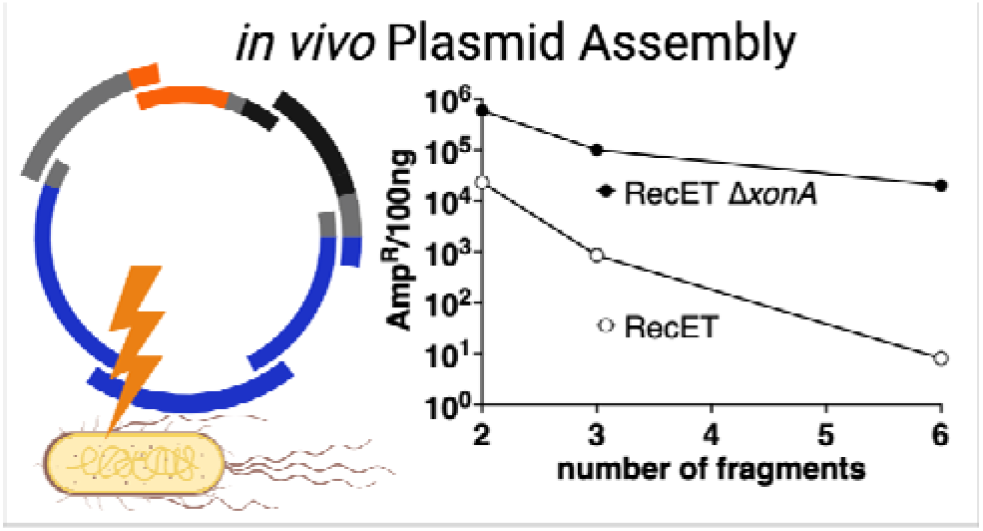

## Notes

### Competing Interest Statement

The authors have declared no competing interest.

