## Supporting Information for "Enhancement of RecET-mediated *in vivo* linear DNA assembly by a *xonA* mutation"

**Inventory of Supporting Information**

**Supplemental Results and Discussion**

**Effect of increasing terminal homology length on linear DNA assembly**

**Effect of *recJ* and *exoX* mutations on linear DNA assembly**

**Supplemental Methods for experiments described in this paper**

**Preparing linear DNA molecules for use in assembly**

**General procedures for linear DNA assembly**

**Table S1. Comparison of values for *in vivo* linear DNA assembly reported by different labs**

**Table S2. *Escherichia coli* K-12 strains and plasmids**

**Table S3. Supplemental *Escherichia coli* K-12 strains**

**Figure S1. Assembly of two linear DNAs with various homology lengths into intact plasmids by the λ Red and RecET systems**

**Figure S2. DNA fragments used for the three-way assembly of pLT59**

**Figure S3. Sequence analysis of white colonies from the pBR-*lacZ* assembly reactions**

**Figure S4. Additional data for *in vivo* linear assembly from six linear dsDNA fragments**

**Supplemental References**

**Supplemental Results and Discussion**

**Effect of increasing terminal homology length on linear DNA assembly.** We found that increasing the length of terminal homology increased the frequency of λ Red-mediated recombination such that with 200bp of homology, Red-mediated recombination is only 10-fold reduced from that of RecET and at >330bp of homology, the difference between the systems is only 5-fold (Figure S1). We also found that the increased recombination frequency observed with longer homologies is not dependent on *E. coli* recombination systems, since mutation of *recA* had no effect on the frequency. The enhancement of Red recombination for linear DNA assembly seen with longer homologies is consistent with our previous results^1^ which tested *in vivo* Red- and RecET-mediated intramolecular circularization of a linear dimer plasmid to form a circular monomeric product. This reaction provided extremely long terminal homologies, with a final circular product of ~4.4 kb and an initial linear substrate of twice that length. This reaction was extremely efficient for both Red and RecET, and the two systems gave similar frequencies for this recombination. We believe that the requirement of longer homologies for the Red system and differences in plasmid allele inheritance that we observed in the previous experiment^1^ both suggest that the two systems process linear substrates differently. Unfortunately any investigation of those differences is beyond the scope of this letter and we encourage other interested researchers to pursue this question further.

**Effect of *recJ* and *exoX* mutations on linear DNA assembly.** In light of the possibility that host single-strand DNA exonucleases other than ExoI could also process linear DNA introduced by electroporation or formed during the recombination reaction, we asked whether removal of two other *E. coli* single-strand DNA exonucleases affected the frequency of recombinant recovery for assembly of the pBR-*lacZ* plasmid. Strain NC558 is deleted for *recJ*, which encodes the RecJ protein, a processive 5'→3' ssDNA exonuclease^2^. When NC558 was tested for assembly of six fragments, we observed an average frequency of 1x10^2^Amp^R^/10^8^ viable cells, similar to that seen in LT1795, the *xonA*^+^ strain, indicating that the *recJ* mutation had no effect. 80% of the recombinant colonies obtained from the *recJ* mutant cells were blue on X-gal indicator agar. 25 white colonies (indicating defects in the *lacZ* gene) were analyzed by isolating their plasmid DNA. Restriction analysis demonstrated that nearly half (11/25) of the *lacZ* mutant plasmids were not assembled correctly in the *recJ* mutant background, a similar percentage to that seen for LT1795, the *recJ^+^* strain.

*E. coli* ExoX is a distributive 3'→5' exonuclease active on both ss- and dsDNA^3^. A strain deleted for the *exoX* gene, NC567, was tested for the ability to assemble six linear DNAs. The average recombination frequency observed with this strain was 5x10^1^Amp^R^/10^8^ viable cells, again similar to the both the wildtype LT1795 strain and the *recJ* mutant. A total of 9 Amp^R^ colonies were recovered and found to be blue on X-gal indicator agar. The *exoX* and *recJ* mutant results suggest that neither of these exonucleases plays a role in the fragment assembly reaction.

**Supplemental Methods for experiments described in this paper.**

**Preparing linear DNA molecules for use in assembly.** When plasmid DNA was used for PCR, ~100 ng of the plasmid was first digested with two different restriction enzymes, producing a linear DNA. The digest preparation was diluted 1000-fold and used as a template for PCR with primers that created fragments with 50bp terminal overlap (unless otherwise noted). In all cases, individual DNA fragments generated by PCR were tested prior to use in DNA assembly by introduction into competent DH5α cells by electroporation. These control transformations were plated on L+Amp agar. An absence of Amp^R^ colonies indicates an absence of any residual uncut plasmid DNA (i.e. confirmed complete digestion) from that used as template.

Linearized pLT59 was used as the PCR template for all fragments used in 2-way and 3-way recombination experiments. The linear DNAs generated with PCR contain a partial (nonfunctional) *ori* and *kan* gene, thus, only precise joining *in vivo* will result in an *ori^+^* recombinant that can replicate and confer kanamycin (Kan) resistance. Assembly of pLT59 from two DNA fragments (1548 and 2490 bp in length) is shown in Figure 1B, assembly of the same plasmid from three DNA fragments (1296, 1327 and 1465 bp in length) is shown in Figure S2.

The same protocol was used to prepare six linear DNAs for recombination from plasmid pLT61 (see Figure S3). This plasmid contains a chloramphenicol resistance cassette, *cat*, immediately flanked by its own promoter and transcription terminator inserted into the *kan* coding region. In this 6-way linear assembly the *ori*, *bla* and *cat* functional regions are each brought in by two separate PCRs. The linear DNAs used for assembly were 941, 766, 912, 875, 902 and 858 bp in length.

We designed the plasmid designated pBR-*lacZ* to further test assembly of six fragments. In pBR-*lacZ*, the tetracycline resistance gene, *tet*, of pBR322 is replaced with the *E. coli* *lacZ* gene, retaining the *tet* regulatory elements (see Figure 5A). To assemble the pBR-*lacZ* plasmid backbone, three separate PCR products were generated, using linearized pBR322 as a template. Both the plasmid *ori* and the *bla* gene are split between two fragments. All three fragments were electroporated separately into cells to ensure that no intact pBR322 plasmid was present. The 3075bp *lacZ* gene is assembled from three separate linear DNA fragments. These DNAs were either generated by colony PCR or synthesized as gBlocks (IDT). All six fragments must be accurately and precisely joined in correct order to result in a plasmid with an intact (a) *ori*, (b) *bla* gene and (c) *lacZ* gene. Amp resistant colonies (Amp^R^) will only be recovered if both the *ori* and *bla* gene are functional. The *lacZ* gene encoding β-galactosidase allows a blue/white colony screen for fidelity of DNA joining of the *lacZ* fragments, only blue colonies express a functional *lacZ* gene. Fragment sizes were 1102, 1030, 1154, 1030, 1040, and 922bp in length.

To create linear DNA fragments of decreasing homology lengths, the oligos used to amplify the linear DNA by PCR were shortened the appropriate number of bases on their 5' end, thus maintaining the same fragment with a shorter homology region.

**General procedures for linear DNA assembly.** The following guidelines are given for cloning genes of interest into plasmid backbones. Although expression vectors with multi-cloning sites are useful for this, any plasmid backbone can be used. See Thomason *et al.*^4^ for detailed recombineering methods.

*Care of strain NC553:* The bacterial strain NC553 is temperature-sensitive since it contains a defective λ prophage with the *recET* and λ *gam* recombination genes under control of the *c*I*857* gene that encodes a temperature-sensitive repressor. Cells must always be grown at 30-32°C. Propagation at higher temperatures will yield poor growth and give rise to mutants unable to express the recombination functions. Upon receiving the strain, a glycerol stock should be made from bacteria grown on either LB solid agar or in LB broth. When growing cells for fragment joining, always grow the strain at 30-32°C until the 42°C heat induction step for expression of the recombination functions. NC553 contains no plasmid or drug markers, making it possible to assemble plasmids with different DNA replication origins and select for a wide range of drug resistances in this background.

*Preparation of vector:* Digest 100ng of the vector, preferably with two unique restriction enzymes (made easier with a multicloning site) to completely linearize the vector. Clean the digest using a commercial kit (such as Qiagen) eluting in 50ul of dH_2_O or elution buffer supplied with the kit. Overhangs left by restriction enzymes are not a concern, and like others ^5^ we have found that target homologies of 10 bases or more internal to the cut site still yield high frequencies of correct fragment joining, with terminal non-homologies removed during the recombination process. If there are no appropriate restriction sites it is possible to PCR amplify the plasmid origin of choice.

In all cases, it is critical that the linear vector preparation be tested to ensure complete restriction enzyme digestion by introducing 1μl of the purified prep into electrocompetent cells (such as commercial DH5α) by electroporation. After a 2 hour outgrowth, spread 100μl cells on a solid agar LB plate containing an appropriate concentration of drug corresponding to the antibiotic resistance marker on the plasmid backbone. The resulting colony count should be <10 colony forming units (cfu). If large numbers of drug resistant cfu are found, plasmid restriction must be improved until background count is lowered; an alternative is to purify the linear fragment from an agarose gel. Do not omit this control of testing the vector alone for complete linearization, since incomplete cutting will give rise to a high background of colonies containing only the vector which will swamp out the yield of assembled plasmids containing your desired fragment(s). Note that simply visualizing the digested vector on an agarose gel to look for supercoiled plasmid is not an adequate control. Trace amounts of uncut plasmid may be invisible on the gel but can still give substantial numbers of drug resistant colonies after transformation.

*Preparation of fragments:* Other DNA fragments to be incorporated onto the plasmid backbone should be designed to contain appropriate 30-50bp homologies at each end. We routinely use 50bp of overlapping homology. Colony PCR ^4^ can be used to amplify genes from the *E. coli* chromosome or a BAC. A plasmid can be used as a template if care is taken to completely linearize it, especially if it contains the same drug resistance you will be selecting for in your assembly product. As with the vector, these DNAs should be purified to remove salts and visualized by agarose gel electrophoresis to insure the correct size fragments. It is not always necessary to quantitate and dilute this DNA as long as adequate yields are obtain, we routinely simply use 1μl of the cleaned PCR fragment. Synthetic dsDNA (i.e. gBlocks from IDT) are reconstituted at 100ng/μl, and 1μl used per recombination.

*Growth and induction:* Details of cell preparation for electroporation and recombinant proficient cells is given in Thomason et al^4^. Briefly, an overnight culture is diluted into fresh L broth and grown at 32°C to log phase. At this point the culture is divided and half of the cells are shifted to 42°C for 15min to induce expression of the *recET* genes, with the remaining culture remaining at 32°C. This 42°C temperature induction requires a shaking water bath in order for efficient heat transfer (see https://redrecombineering.ncifcrf.gov/). The short induction time is critical as prolonged expression of the recombinations functions is toxic. After 15min both flasks are rapidly chilled in an ice water slurry. The cells are then made electrocompetent. We have not tried using calcium-competent cells for linear DNA assembly.

*Electroporation and outgrowth:* The DNAs to be assembled are introduced into electrocompetent and RecET recombination proficient (induced at 42°C) cells in which recombinants are generated. Uninduced (32°C) cells are used as a control to check for the presence of uncut vector^4^. After a 2hr outgrowth in 1ml LB broth at low temperature (30-32°C), five consecutive 10-fold serial dilutions of the culture are made into sterile M9 salts or Tris-Magnesium (TM) ^4^. From the last two dilutions (10^-4^ and 10^-5^), 100μl of the mixture is spread on LB agar to determine total viable cell counts. From the undiluted culture and the 10^-1^ and 10^-2^ dilutions, 100μl is spread on LB agar containing the appropriate drug to select for intact plasmids. Plates are incubated at 30-32°C overnight and scored on LB agar with the appropriate drug for bacterial colonies. Few if any colonies should be found with the uninduced 32°C cultures. Such drug resistant colonies from the uninduced culture indicate uncut plasmid background. The recombinant frequency is calculated using titers from the induced (42°C) RecET plates as: (drug resistant colonies/total viable colonies)10^8^.

*Identifying recombinant clones:* Purify 12 colonies picked from the drug plates for the induced 42°C cultures onto the same solid medium and incubate these plates at 30-32^o^C. Incubating the purification plates at higher temperatures will induce the RecET recombination functions and kill the cells. These purified isolates can be screened for plasmid assembly either with colony PCR or by isolating DNA using a miniprep. For the former approach, perform colony PCR ^4^ on the 12 purified isolates to confirm recombinant junctions and screen for desired insertion/s. The PCR primers should be designed to span unique junctions joined together during assembly, to confirm assembly of the linear DNAs into plasmids. For smaller inserts, primers can be designed to span the entire region. These PCR products can then be directly sequenced. Alternatively, the purified colonies can be used to start overnight cultures for plasmid DNA minipreps for further analysis by restriction enzyme digestion, agarose gel analysis and sequencing. We strongly recommend that all insertions be sequenced in their entirety, as well as recombinant junctions, to ensure that correct clones are identified. As noted in this paper, mistakes in inserts usually arise from either PCR or gBlock synthesis. We also note that Lac^+^ (blue) isolates were sequenced and even though the LacZ protein was functional, mutations resulting in amino acid changes were found. Of the seven Lac^+^ plasmids sequenced from PCR assembly, all contained at least one mutation. Seven Lac^+^ plasmids from gBlock assembly were also sequenced, four of these contained at least one mutation. On some occasions we have seen frameshifts occurring at the beginning of an open reading frame when the gene to be cloned conferred toxicity on the bacterial cells.

**Table S1. Comparison of values for *in vivo* linear DNA assembly reported by different labs**

**Average DrugR per ~100ng vector***

| **# Fragments** | **RecET ∆*xonA*^1^** | **RecET^2^** | **iVEC^3^** | **ZeBrα^4^** | **ExoCET^5^** | **Gibson^5^** | **Gibson+ RecET^5^** |
| --- | --- | --- | --- | --- | --- | --- | --- |
| **2** | **5.9x10^5^** | **2.8x10^5^** | **4.3x10^3^** | **3.7x10^3^** |  |  |  |
| **3** | **1.0x10^5^** |  | **2.1x10^2^** | **7x10^3^** |  |  |  |
| **4** |  | **~1x10^4^** | **1.9x10^2^** | **5x10^2^** |  |  |  |
| **5** |  | **~7x10^1^** |  | **9x10^2^** |  |  |  |
| **6** | **2.0x10^4^** |  |  |  |  |  |  |
| **7** |  |  | **4.7x10^1^** |  | **1.5x10^2^** | **1.4x10^4^** | **3.0x10^4^** |
| **13** |  |  |  |  | **7.3x10^1^** | **5x10^2^** | **1.1x10^3^** |

^*^ Values represent approximate colony counts and were calculated by normalizing to ~100ng of the fragment that contained the selected drug resistance. In most cases, all fragments were used at the same DNA concentration.

^1^ Present results.

^2^ Fu *et al*. ^5^ Figure 1D and supplemental Figure 1 for two fragments. For four fragments, Baker *et al.* ^6^ Figure 2B, 5 fragments, Supplemental Figure 2B.

^3^ Nozaki and Niki ^7^. From Figure 5B, 30bp homology and three fragments. Figure 5E, 20bp homology and three fragments. Figure 5E, 40bp homology and seven fragments. The authors used a three-fold excess of insert fragments relative to the linear vector backbone.

^4^ Ritcher *et al*. ^8^ From Figure S2, two fragments. From Figure 3C and Figure 4A, three and four fragments. From Figure 8C, five fragments. For three-way assembly, 100ng vector and a three-fold molar excess of inserts were used.

^5^ Song *et al.* ^9^ From Figure 3B and 3D. Six candidates were analyzed for the 7 fragment reaction, and twelve for the 13 fragment reaction. All plasmids assembled from 7 fragment with ExoCET were correct by restriction analysis, while ~half of the plasmids assembled from 13 fragments were correct by this criteria. None of the candidates assembled with Gibson alone were correct. A quarter to a third of the plasmids formed with the combined Gibson-RecET method were correct, the rest had fragments omitted. Thus, the increase in frequency achieved by including Gibson assembly came at the expense of reduced accuracy.

**Table S2. *Escherichia coli* K-12 strains and plasmids**

| **Strain name** | **Relevant genotype** | **Source** |
| --- | --- | --- |
| HME6 | W3110 Δ*lacU169 galK_TYR145UAG_* λ*c*I*857* Δ*(cro-bioA)* | Ellis *et al.*^10^ |
| JS663 | W3110 Δ*lacU169 galK490* [λ *cI857* (*int*-*cIII*<>*cat exo bet* Δ*gam*) Δ (*cro-bioA*)] | this study |
| LT1795 | W3110 Δ*lacU169 galKTYR145UAG* λ*cI857*Δ (*cro-bioA*) [λ (*int-cIII*)< >*gam* *recE recT*] | this study |
| NC540 | W3110 Δ*lacU169 galKTYR145UAG* λ*cI857*Δ(*cro-bioA*) Δ*xonA* | this study |
| NC553 | W3110 Δ*lacU169 galKTYR145UAG* Δ*xonA* λ*cI857* Δ(*cro-bioA*) (*int-cIII*)< >*gam* *recE recT* | this study |
| SIMD63 | W3110 Δ*lacU169 galKTYR145UAG* λ*cI857* Δ (*cro-bioA*) [λ (*int-cIII*)< >*recE recT*] | Thomason *et al.*^4^ |
| Plasmid name | Relevant genotype | Source |
| pLT59 | pUC plasmid *bla^+^ kan^+^* | Thomason *et al.*^11^ |

**Table S3. Supplemental *Escherichia coli* K-12 strains**

| **Strain Name** | **Relevant genotype** | **Source** |
| --- | --- | --- |
| JS677  JS679  JS726  JS727  NC558  NC567 | JS663[pSIM26] Tet^R^ (Gam producing plasmid)  W3110 Δ*lacU169 galKTYR145UAG* [λ *cI857 recE recT* Δ(*cro-bioA*)] [pSIM26] Tet^R^  JS679 *recA*<>*spec*  JS677 *recA*<>*spec*  W3110 Δ*lacU169 galKTYR145UAG* Δ*recJ<>cat* λ*cI857* Δ(*cro-bioA*) (*int-cIII*)< >*gam* *recE recT*  W3110 Δ*lacU169 galKTYR145UAG* Δ*exoX<>spec* λ*cI857* Δ(*cro-bioA*) (*int-cIII*)< >*gam* *recE recT* | this study  this study  this study  this study  this study  this study |
| Plasmid name | Relevant genotype | Source |
| pLT61 | pUC plasmid *bla^+^ cat^+^* | Thomason *et al.*^11^ |

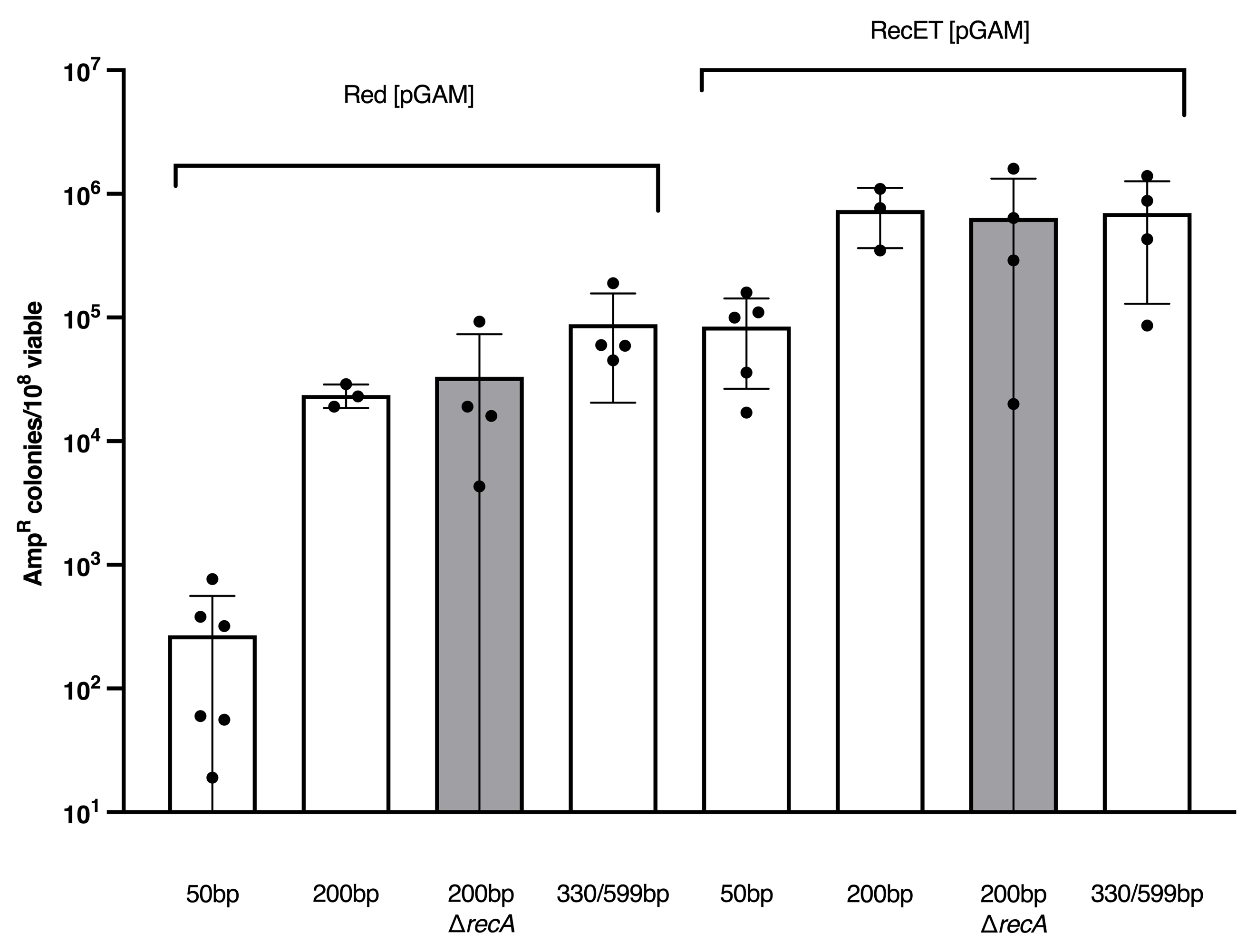

**Figure S1. Assembly of two linear DNAs with various homology lengths into intact plasmids by the λ Red and RecET systems.** Recombinant frequency is expressed as the number of Amp^R^ colonies/10^8^ total colonies for the indicated recombineering systems. In all cases λ Gam protein was supplied from the low-copy tetracycline resistant plasmid pSIM26. Homology lengths are indicated on the x-axis, in one case they were 330bp in the plasmid origin and 599bp in the *kan* gene. In the grey bars, the cells are also mutant for *recA*. Results from independent experiments are indicated by the black circles. Assembly was dependent on supplying both fragments and on expression of the λ Red or RecET functions. Tetracycline was used at 25 μg/ml for plasmid maintenance. Error bars indicate s.d.

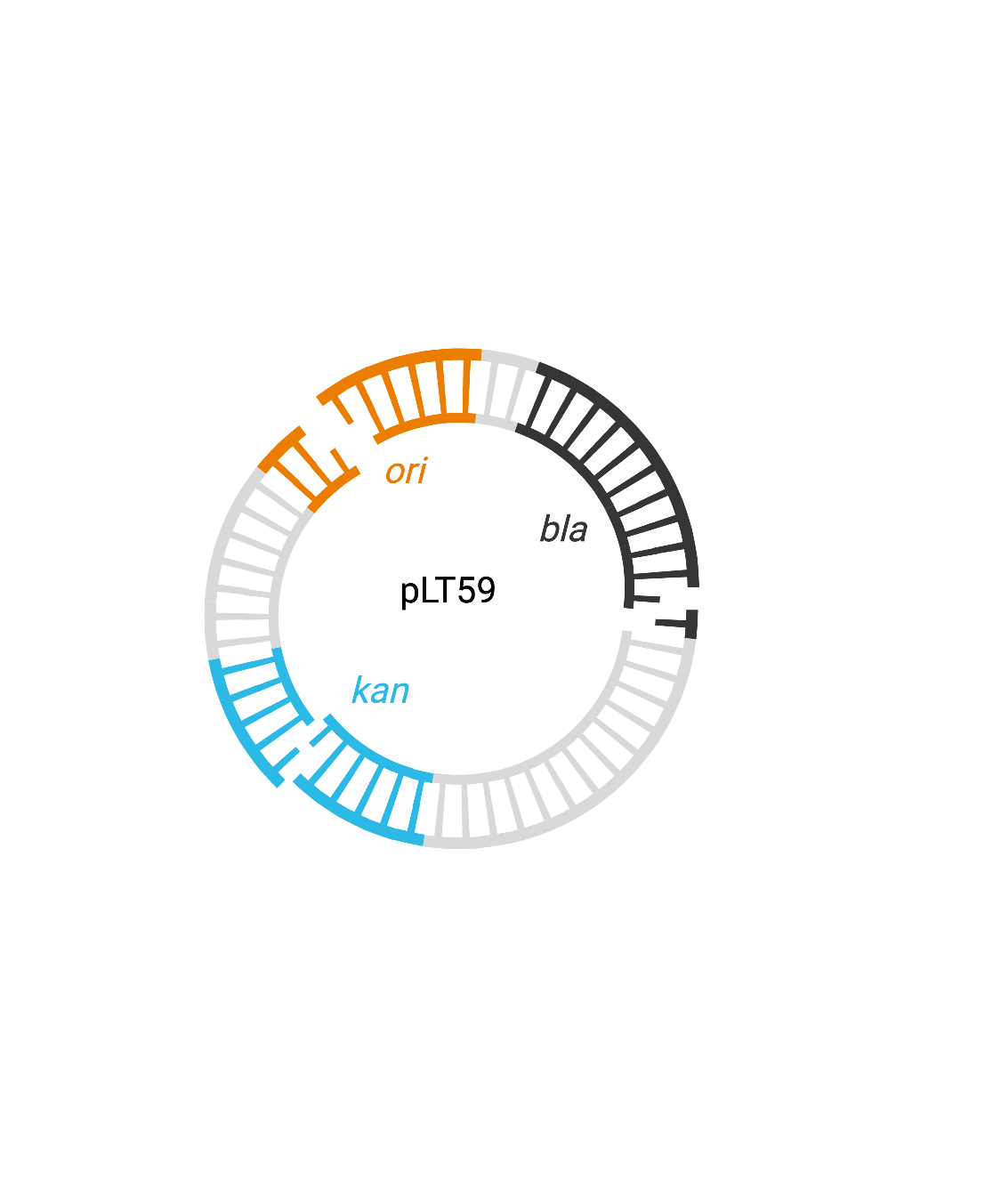

**Figure S2. DNA fragments used for the three-way assembly of pLT59.** Terminal homologies present in each fragment within the *ori*, *bla*, and *kan* genes are indicated by the single-strand bases. One single-strand base indicates a homology of 50-60 bases. The data are shown in Figure 2 of the main paper.

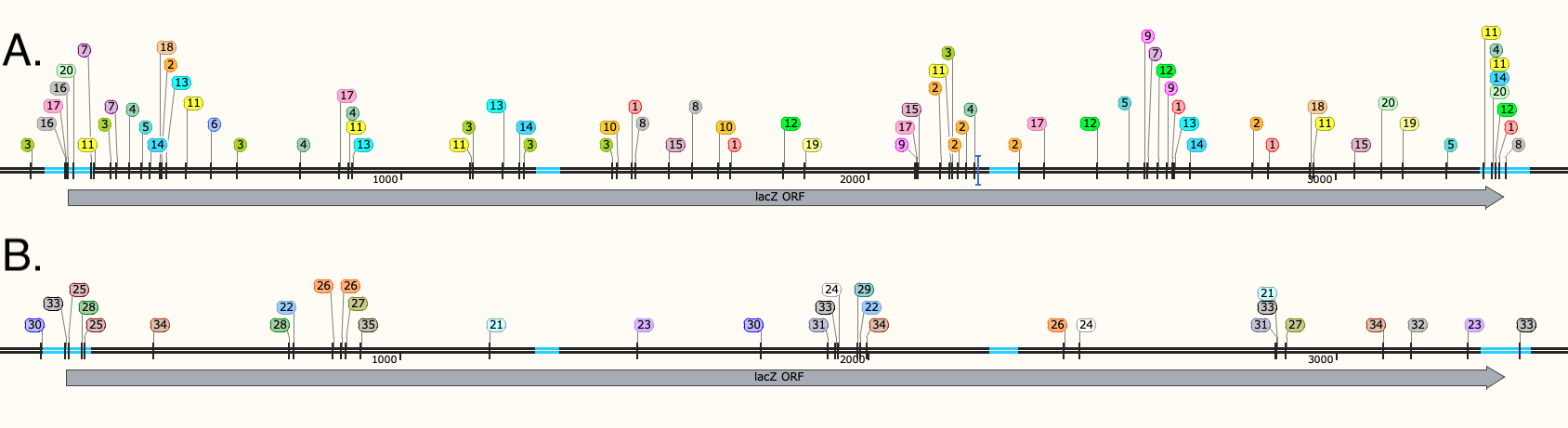

**Figure S3. Sequence analysis of white colonies from the pBR-*lacZ* assembly reactions.**

The black bar with cyan inserts represents a portion of the pBR-*lacZ* plasmid. The homology overlaps between fragments used for assembly are shown in cyan. The 3075bp *lacZ* open reading frame *(orf)* is indicated by the grey arrow. The region from the promoter to the translational stop was sequenced; locations of observed mutations are indicated with vertical lines and a colored circle with a number, indicating the isolate number. If multiple mutations were found in an isolate, they are indicated with the same color and number. We did not find increased numbers of mutations in the overlapping homology region where the single-strand annealing occurs A. PCR products were used for assembly. A total of 20 isolates were sequenced and among these, 68 point mutations and 6 insertion or deletions were found. 9 candidates had mutations in the overlap homologies. B. gBlocks were used for assembly. A total of 20 isolates were sequenced and among these, 17 point mutations and 14 insertion or deletions were found. Only 3 isolates had any mutations in the overlap homologies. The 5 candidates not shown were isolates that had apparent synthesis errors containing small repeats.

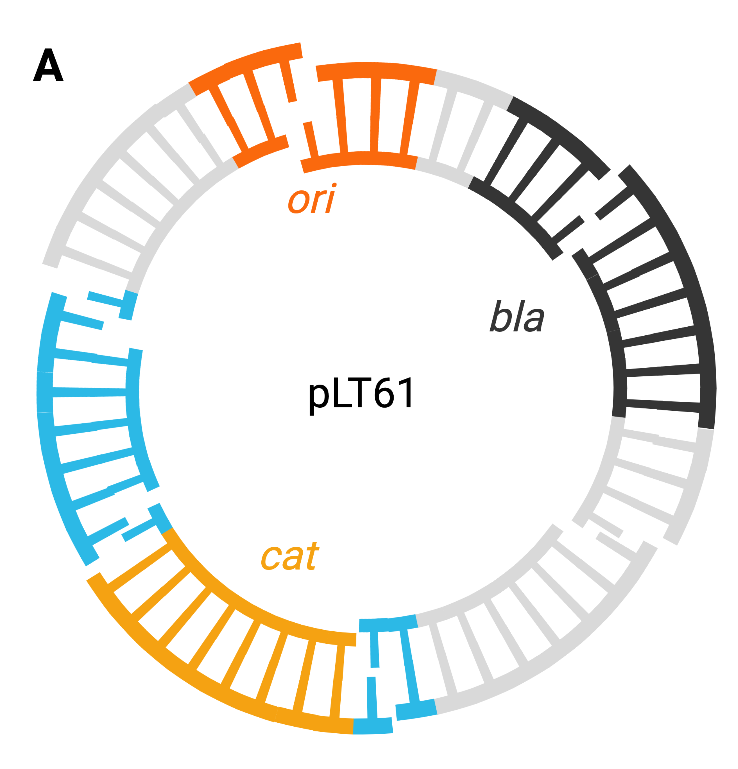

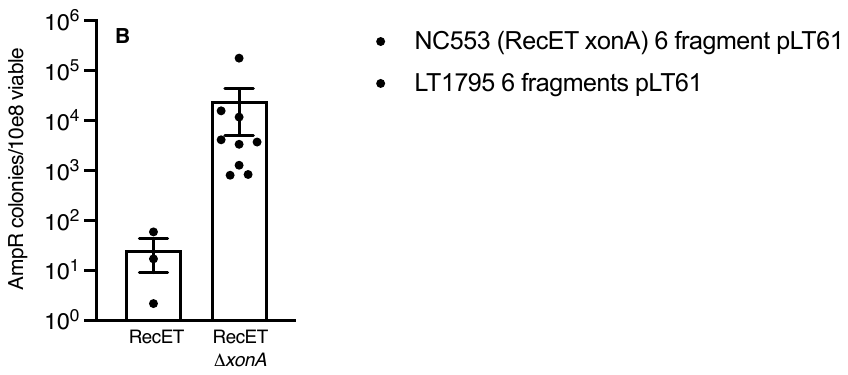

**Figure S4. Additional data for *in vivo* linear assembly from six linear dsDNA fragments. A.** Six PCR fragments were used to assemble the circular plasmid pLT61. Of the six junctions, one is within the plasmid *ori*, a second is within the *bla* gene, and two more are within the chloramphenicol resistance gene, *cat*. **B.** Elimination of the host ExoI function (i.e. Δ*xonA*) increased the frequency of plasmid assembly ~1000 fold. When Amp^R^ colonies from the RecET Δ*xonA* recombination were scored for Cm^R^, 194/200 colonies were Cm^R^, indicating accurate joining of the linear DNA segments. We anticipate that most of the Cm^S^ plasmids arose from PCR mistakes or primer synthesis mistakes, as found for the *lacZ* gene in other six-way assembly experiments (see Figure 5, main paper).
